# 3-5 Hz membrane potential oscillations decrease the gain of neurons in visual cortex

**DOI:** 10.1101/069252

**Authors:** Michael C. Einstein, Pierre-Olivier Polack, Peyman Golshani

## Abstract

Gain modulation is a computational mechanism critical for sensory processing. Yet, the cellular mechanisms that decrease the gain of cortical neurons are unclear. To test if low frequency subthreshold oscillations could reduce neuronal gain during wakefulness, we measured the membrane potential of primary visual cortex (V1) layer 2/3 excitatory, parvalbumin-positive (PV+), and somatostatin-positive (SOM+) neurons in awake mice during passive visual stimulation and sensory discrimination tasks. We found prominent 3-5 Hz membrane potential oscillations that reduced the gain of excitatory neurons but not the gain of PV+ and SOM+ interneurons, which oscillated synchronously with excitatory neurons and fired strongly at the peak of depolarizations. 3-5 Hz oscillation prevalence and timing were strongly modulated by visual input and the animal’s behavior al response, suggesting that these oscillations are triggered to adjust sensory responses for specific behavioral contexts. Therefore, these findings reveal a novel gain reduction mechanism that adapts sensory processing to behavior.

## INTRODUCTION

Gain modulation is a fundamental mechanism by which the brain adjusts the strength of sensory signals (Salinas & Sejnowski, 2001). During behavior, neuronal gain is tuned moment -by-moment in order to prioritize information streams important for meeting immediate behavioral demands (Harris & Thiele, 2011; Posner 1980). Notably, attention has been found to either increase (Moran & Desimone, 1985; Motter, 1993; Roelfsema et al., 1998; Chalk et al., 2010) or decrease (Luck et al., 1997; Reynolds et al., 1999; Treue & Maunsell, 1996) the gain of neurons throughout the visual cortex to prioritize coding and perception of attended cues.

Several cellular and network mechanisms that increase the gain of sensory neurons during behavior have already been identified. Signals from the prefrontal cortex (Zhang et al., 2014; Gregoriou et al., 2014; Moore & Armstrong et al., 2003), thalamus (McAlonan et al., 2008; Purushothaman et al.,2012; Wimmer et al., 2015), and neuromodulatory centers (Polack et al., 2013; Pinto et al., 2013; Fu et al., 2014) have all been shown to increase the gain of visual cortical neurons in behaving animals. However, mechanisms that reduce the gain of sensory cortical neurons during behavior are still poorly understood. Recruitment of inhibitory GABAergic interneurons has been implicated as a mechanism that could reduce the gain of visual and auditory cortical neurons in behaving animals (Katzner et al., 2011; Disney et al., 2007; Soma et al., 2012; Olsen et al., 2012; Schneider et al., 2014). Yet, the cellular mechanisms that decrease neuronal gain in sensory cortices during behavior remain unclear.

We hypothesized that low frequency subthreshold oscillations could be a mechanism that reduces neuronal gain during behavior. Previously associated with sleeping and anesthetized states (Steriade et al., 1993), low frequency subthreshold oscillations have recently been observed in rodent visual (Polack et al., 2013; Bennet et al., 2013), barrel (Poulet & Petersen, 2008), auditory (Zhou et al., 2014; Schneider et al., 2014) and motor (Zagha et al., 2015) cortex neurons of awake behaving animals. During low frequency oscillations, neurons’ baseline membrane potential was significantly hyperpolarized (Zagha et al., 2015; Bennet et al., 2013), which could decrease the responsiveness of neurons to incoming signals (Cardin et al., 2008; Carandini & Ferster, 1997; Nowak et al., 2005). Moreover *, in vivo* (Cohen & Maunsell, 2009; Fries et al., 2001) and *in vitro* (Volgushev et al., 1998; Lampl & Yarom, 1993) experiments suggest that low frequency oscillations could provide timing templates that filter inbound sensory signals of a different time structure, which could effectively reduce the gain of sensory cortex neurons (Engel et al., 2001; Schroeder & Lakatos, 2009).

To investigate this hypothesis, we performed whole-cell recordings of V1 L2/3 excitatory, parvalbumin-positive (PV+), and somatostatin-positive (SOM+) neurons in awake and behaving animals. We found prominent low frequency (3-5 Hz) membrane potential oscillations in all neuron types. These 3-5 Hz oscillations decreased the spontaneous firing rate and gain of excitatory neurons. Meanwhile, PV+ and SOM+ interneurons oscillated in phase with excitatory neurons, but fired strongly at the depolarized peaks of these oscillations. 3-5 Hz oscillation recruitment depended on both visual processing and behavioral state. Visual stimulation significantly increased the prevalence of oscillations, and engagement on a visual discrimination task strongly influenced the initiation, duration, and prevalence of oscillations. Altogether, our findings suggest that 3-5 Hz subthreshold oscillations are a novel mechanism for decreasing neuronal gain to tune sensory processing according to an animal’s specific behavioral context.

## RESULTS

### 3-5 Hz Vm oscillations are highly stereotyped events that reduce the gain excitatory neurons

We performed two-photon guided whole-cell Vm recordings from 40 excitatory, 6 PV+, and 7 SOM+ L2/3 V1 neurons in head-fixed mice free to run or rest on a spherical treadmill (Figure 1A, B). For each recording, electrocorticogram (ECoG) activity was simultaneously acquired within the vicinity (300-500 μm) of the patch-clamp pipette tip was simultaneously acquired. In all our recordings, we detected epochs of high amplitude 3-5 Hz Vm oscillations (Figure 1A) that typically lasted for 1-2 seconds (1.6 ± 0.05 seconds, n = 53; Figure 1C, D). During oscillatory events, the neuron’s baseline Vm substantially hyperpolarized (Mean = -12.0 ± .61 mV, n = 53) and displayed high amplitude (>10 mV) rhythmic depolarizations at 4.14 ± 0.06 Hz (n = 53; range = [2.94, 5.04]). The oscillation frequency, duration, and baseline hyperpolarization were similar in excitatory, PV+, and SOM+ neurons ( one-way ANOVA p = 0.55; Figure 1D). However, PV+ interneurons exhibited larger amplitude depolarizing events ( one-way ANOVA, p= 0.01) than excitatory (Tukey-HSD, p=0.04) and SOM+ (Tukey-HSD, p = 0.01) neurons did during oscillatory periods. The mean firing rates of excitatory neurons significantly decreased during the oscillation, with excitatory neurons rarely firing action potentials during the oscillatory episodes (Spontaneous firing rate- No oscillation: 1.34 ± .05 Sp.s^-1^, Oscillation: 0.55 ± .02 Sp.s^-1^; WSRT, p = 0.002; Figure 1B, 2C). In contrast, PV+ and SOM+ interneurons still fired strongly at the peaks of oscillations (13.3 ± 1.03 Sp.s^-1^, n=6, and 6.23 ± 0.8 Sp.s^-1^, n=7, respectively; Figure 1B, 2C).

**Figure 1.**
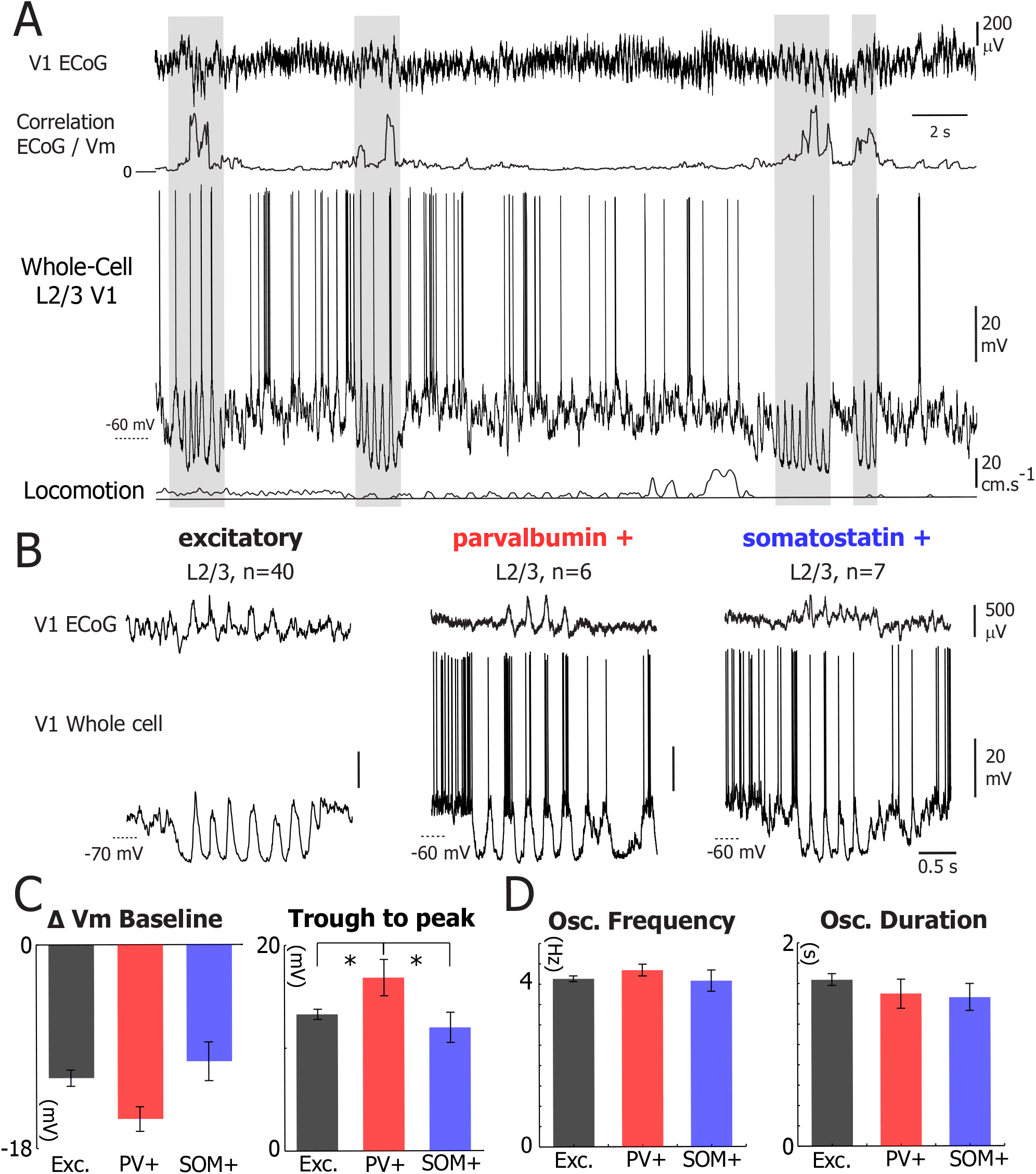
V1 L2/3 excitatory, PV+, and SOM+ neurons’ Vm spontaneously undergo high amplitude 3-5Hz oscillations. (A) Example whole cell recording from a V1 layer 2/3 excitatory neuron during wakefulness, simultaneously recorded with the local electroencephalogram (ECoG, top) and the treadmill motion (locomotion, bottom). The second trace from the top represents the correlation between the ECoG and the membrane potential (Vm) measured with the whole-cell recording. Grey highlights indicate times when 3-5 Hz oscillations were observed i n the neuron’s Vm. (B) Simultaneous V1 ECoG (top) and whole-cell recordings (bottom) from V1 L2/3 excitatory (left), PV+ (center), and SOM+ (right) neurons during Vm 3-5 Hz oscillations. During Vm 3-5 Hz oscillations, the ECoG also displays prominent 3-5 Hz oscillations. (C) Plots of the mean change in Vm baseline during 3-5 Hz oscillations (left) and mean oscillation trough to peak amplitude (right) for excitatory (black, n=40), PV+ (red, n=6), and SOM+ (blue, n=7) neurons. Error bars represent SEM. PV neurons exp erienced greater changes in trough to peak amplitude (one-way ANOVA, p = 0.01) than excitatory neurons (Tukey-HSD, p = 0.01) and SOM+ neurons (Tukey-HSD, p = 0.04) during Vm 3-5 Hz oscillations. Change in Vm baseline was unchanged between neuronal types (one-way ANOVA, p = 0.10). (D) Plots of mean frequency (left) and duration (right) of 3-5 Hz oscillatory periods in excitatory (black, n=40), PV+ (red, n = 6), and SOM+ (blue, n = 7). Error bars represent SEM. Oscillation frequency and duration was unchanged between neuronal types (one-way ANOVA, p = 0.55 & p = 0.43, respectively).

**Figure 2.**
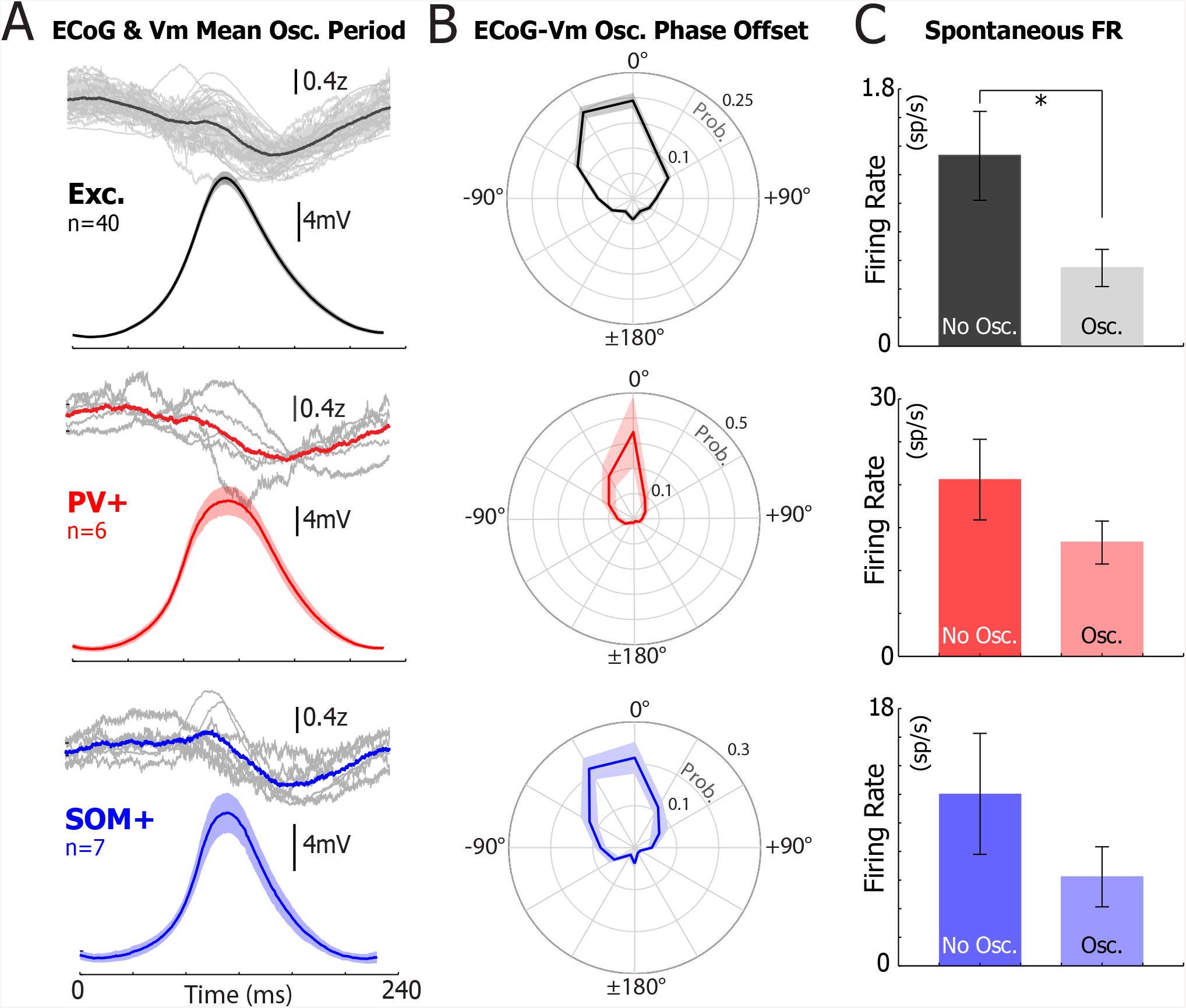
Vm 3-5 Hz oscillations occur synchronously in V1 L2/3 neurons and decrease spontaneous excitatory neuronal output. (A) The mean ECoG (top) and Vm (bottom) during a single period of a Vm 3-5 Hz oscillation for excitatory (black, n=40), PV+ (red, n=6), and SOM+ (blue, n=7) neurons. For the ECoG traces, the colored line represents the mean ECoG z-score of all the neurons, and each light gray trace is the mean ECoG z-score trace from an individual neuron. For the Vm traces, the colored line represents the mean Vm from all cells, and the shaded region represents ±SEM. (B) The mean ECoG-Vm phase offset histogram between 3-5 Hz oscillations detected simultaneously in the ECoG and Vm traces for excitatory (black, n=40), PV+ (red, n=6), and SOM+ (blue, n=7) neurons. The dark line represents the mean phase offset in degrees between the ECoG and the Vm, and the shaded region represents ±SEM. (C) The mean spontaneous firing rate of excitatory (black, n=40), PV+ (red, n=6), SOM+ (n=7) during periods without (no osc.) and with (osc.) Vm 3-5 Hz oscillations. 3-5 Hz oscillations significantly reduced the spontaneous firing rate of excitatory (WSRT, p=0.002) but not PV+ neurons (WSRT, p=0.13) and SOM+ neurons (WSRT, p=0.25).

3-5 Hz Vm oscillations were associated with prominent fluctuations (~500 μV) in the simultaneously recorded ECoG (Figure 1A). The correlation coefficient between Vm and ECoG recordings increased during 3-5 Hz Vm oscillations from 0.002 ± 0.006 to 0.21 ± 0.03 (n=53, WSRT, p= 1.5 x 10^-6^; Figure 1A,1B, 2A). Given the Vm and ECoG correlation during 3-5 Hz oscillations and the similar characteristics of 3-5 Hz oscillations in excitatory, PV+, and SOM+ neurons, we hypothesized that 3-5 Hz oscillations occurred synchronously in L2/3 V1 neurons. To test this hypothesis, we measured the mean phase offset between ECoG and Vm and found no differences between excitatory, PV+, and SOM+ neurons (Excitatory neurons: -7.5° ± 2.2°, PV+ neurons: -12.3° ± 3.8, SOM+ neurons: -14.6° ± 3.1; one-way ANOVA, p = 0.28; Figure 2B). These results suggest that the Vm of excitatory, PV+ and SOM+ neurons excitatory, PV+, and SOM+ neurons oscillated in phase, depolarizing and hyperpolarizing synchronously during each oscillatory cycle.

Because excitatory neurons’ alternate during 3-5 Hz oscillations between hyperpolarized periods and depolarized phases where they likely receive strong inhibitory inputs, we hypothesized that excitatory neurons’ gain could decrease during 3-5 Hz oscillations. To investigate this hypothesis, we recorded the Vm from excitatory (n=40), PV+ (n=6), and SOM+ (n=7) neurons while mice were presented with full-screen drifting gratings (Figure 3). In the presence of oscillations, the mean firing rate of excitatory neurons was strongly reduced for the preferred visual stimulus (2.82 ± 0.71 Sp.s^-1^ No Osc.; 0.75 ± 0.18 Sp.s^-1^ Osc.; n=40 neurons; WSRT, p = 8.1 x 10^-5^; Figure 3C). Yet, the mean orientation selectivity index (OSI) of excitatory neurons was unchanged (WSRT, p = .93; see methods for OSI calculation; Figure 3 —figure supplement 1). Oscillations did not change PV+ (WSRT, p= 0.07) and SOM+ (WSRT, p= 0.63) neurons’ response to the visual stimulus that evoked the greatest response (Figure 3C). In all neurons, the mean firing rate evoked by all non-preferred stimuli was not influenced by the oscillations (excitatory, WSRT, p = 0.97; PV+, WSRT, p = 0.15; SOM+, WRST, p = 0.16). As a result, we conclude that 3-5 Hz oscillation epochs selectively reduced the gain of excitatory neurons during passive viewing.

**Figure 3.**
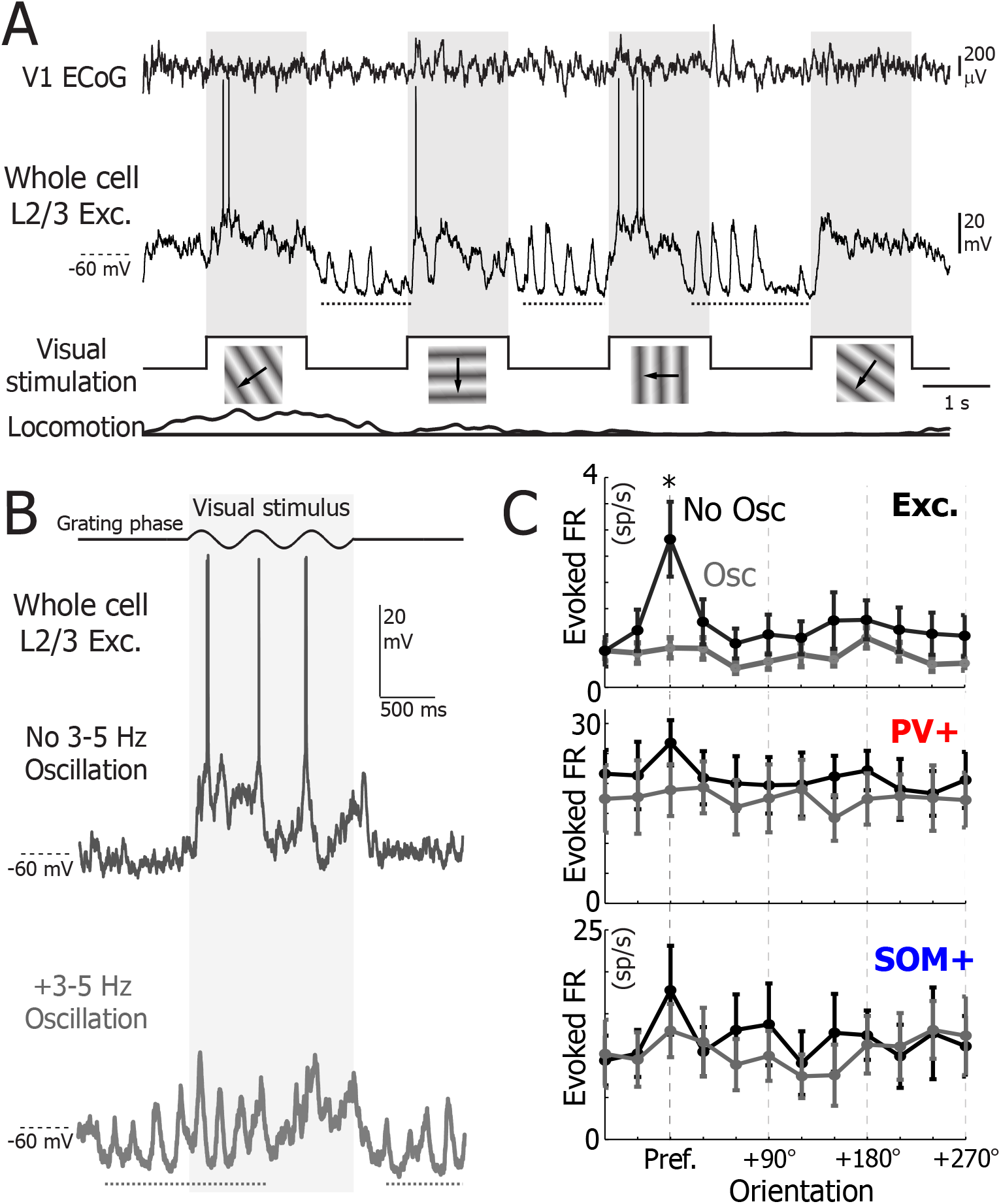
Vm 3-5 Hz oscillations reduce excitatory neuron responsiveness to preferred stimuli during passive viewing of drifting gratings. (A) Simultaneous recordings of the Vm from a layer 2/3 excitatory neuron, local ECoG, visual stimulations, and animal locomotion as an awake animal was shown drifting gratings. Full-field drifting grating presentations lasted 1.5 seconds and were interspersed with 1.5 seconds of an isoluminant gray screen. See Methods for more information about the visual stimuli. Visual stimulus presentation times are highlighted in gray, and dotted lines underline periods of 3-5 Hz oscillations in the Vm recording. (B) Example of an excitatory neuron’s Vm in response to its preferred visual stimulus in the absence (top) and during (bottom) Vm 3-5 Hz oscillations. The dotted lines underline periods of 3-5 Hz oscillations in the Vm recording. (C) The mean orientation tuning of excitatory (top, n=40), PV+ (middle, n=6), SOM+ (bottom, n=7) neurons during (grey) and in the absence of (black) 3-5 Hz oscillations. The firing rate at the preferred angle was significantly larger in the absence of oscillations for excitatory neurons (WRST, p = 8.1x10^-5^), but not for PV+ (WRST, p = 0.07) and SOM+ (WRST, p = 0.63) neurons. Shaded regions indicate ±SEM.

### 3-5 Hz oscillations are more prevalent during passive viewing than during spontaneous activity and occurred at visual stimulus offset

3-5 Hz oscillations were more likely to occur while animals were shown alternations of drifting gratings and grey screens (passive viewing) than during spontaneous activity (defined as periods longer than 5 minutes where animals were shown an isoluminant grey screen; Figure 4A). The incidence rate of oscillations strongly increased in excitatory (WSRT, p = 1.5 × 10^-5^), PV+ (WSRT, p = 0.025), and SOM+ (WSRT, p = 0.038) neurons, during periods of passive visual stimulation compared to periods of spontaneous activity (Figure 4A). There was no difference in 3-5 Hz oscillation incidence between excitatory, PV+, or SOM+ during passive viewing ( one-way ANOVA, p = 0.67) and spontaneous activity (one-way ANOVA, p = 0.38).

**Figure 4.**
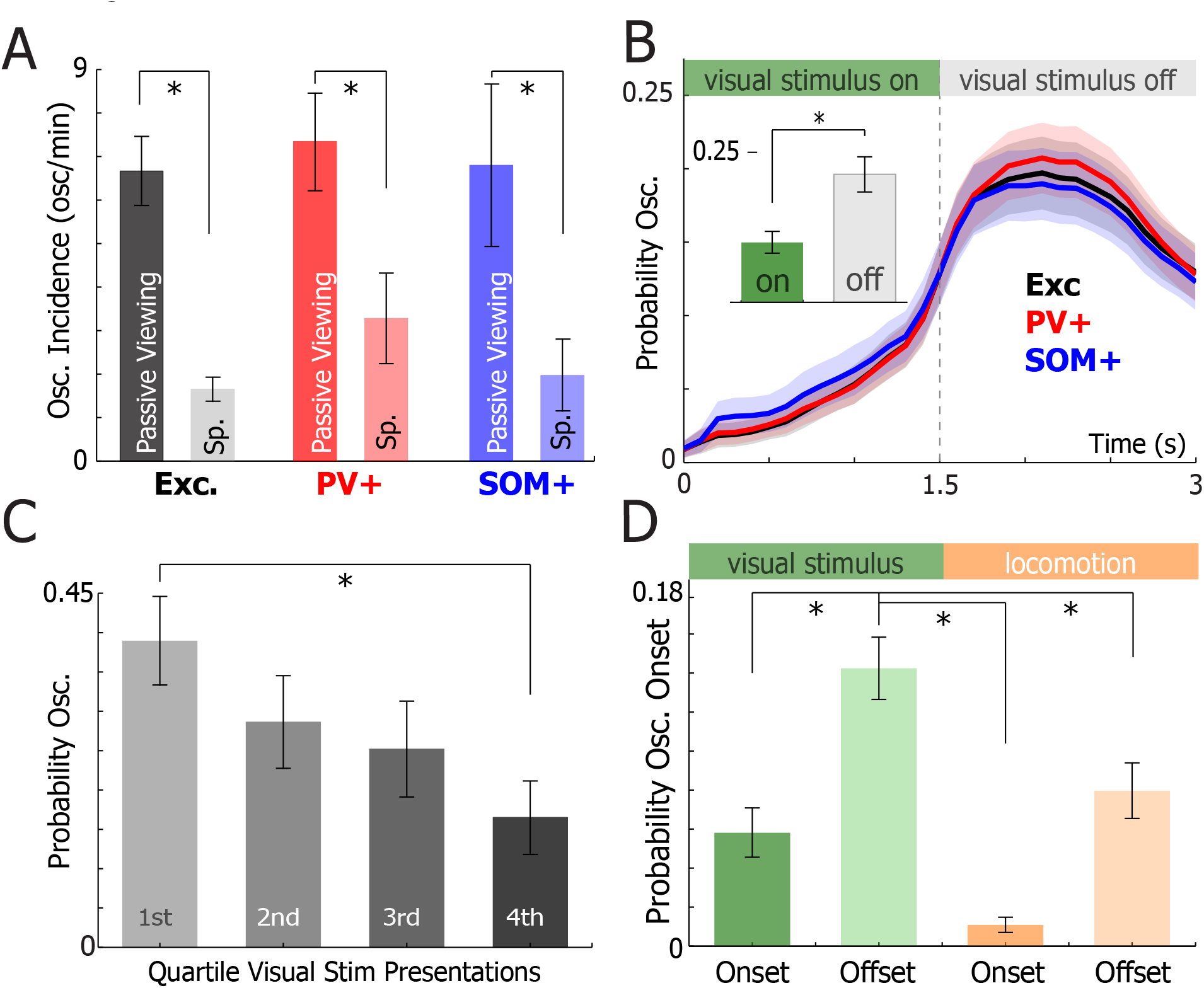
Prevalence and timing of 3-5 Hz oscillations during passive viewing. (A) The number of oscillations per minute during passive viewing (darker) and spontaneo us activity (Sp., lighter) for excitatory (grey), PV+ (red), and SOM+ (blue) neurons. The incidence of oscillations was different for all neuron types during passive viewing and spontaneous activity ( excitatory, WSRT, p = 1.5 x 10^-5^; PV+, WSRT, p = 0.025; SOM+, WSRT, p = 0.038). (B) The mean probability of 3-5 Hz oscillations occurring during and after a visual stimulus for excitatory, PV+, and SOM+ neurons. Shaded regions indicate ±SEM. Inset: the probability of an oscillation occurring for all neuron types w hen a visual stimulus was on and off. Oscillations occurred more frequently after visual stimulus offset than during visual stimuli presentations (WSRT, p = 7.2 x 10^-9^). (C) The mean probability of 3-5 Hz oscillations occurrences was calculated during blocks of visual stimuli presentations grouped by the time of presentation (1^st^quartile = first quarter of visual stimuli shown) for all neurons (n = 31). Recordings with fewer than 100 visual stimulus presentations were excluded (mean number of visual stimuli per neuron = 176 ± 20). The probability of 3-5 Hz oscillations decreased over the course of visual stimulus presentations ( one-way ANOVA, p = 7.7x10^-7^; quartile 1 vs. quartile 4, WSRT Bonferroni Corrected, p = 0.00001). Error bars represent ±SEM. (D) The probability of oscillation onset triggered at visual stimulus (green) and locomotion (tan) onset (colored) and offset (grey). The probability of oscillation initi ation at visual stimulus onset was greater than that at visual stimulus onset, locomotion onset, and locomotion offset (WSRT Bonferroni Corrected, p = 0.024, p = 0.0003, p = 0.003, respectively). Error bars represent ±SEM.

During passive viewing of either 1.5 or 3 second visual stimuli, 3-5 Hz oscillations primarily occurred after visual stimulus offset (Figure 4B and Figure 4—figure supplement 1). In all recorded neurons, the mean probability of 3-5 Hz oscillations following a 1.5 or a 3 second visual stimulus was 2.2 and 2.5 fold greater, respectively, than the probability of 3-5 Hz oscillations occurring during visual stimuli (1.5 s stimuli: n=53, WSRT, p = 7.2 × 10^-9^; 3 s stimuli: n = 9, WSRT, p = 0.004; Figure 4B, Figure 4—figure supplement 1). Interestingly, the probability of an oscillation triggered during or after a passively viewed visual stimulus decreased from the first quartile of visual stimuli to the final quartile of visual stimuli (n = 31 neurons; mean # stimuli presentations per recording = 176 ± 10, repeated measures one-way ANOVA, p = 7.7e-7, WSRT Bonferroni Corrected, p = 0.0001; Figure 4C). As locomotion alters L2/3 V1 neuron Vm dynamics (Polack et al., 2013; Reimer et al., 2014; Bennett et al., 2013), we also analyzed the influence of locomotion on 3-5 Hz oscillation initiation. The probability of oscillation initiation at visual stimulus offset was higher than that at visual stimulus onset, locomotion onset, and locomotion offset (WSRT Bonferroni Corrected, p = 0.024, p = 0.0003, p = 0.003, respectively).

Therefore, synchronized 3-5 Hz oscillations decreased excitatory neuron excitability and were more prevalent when visual stimuli were presented. These findings suggest a role for 3-5 Hz oscillations in modulating visual information processing. Yet, oscillations occurred primarily at the offset of visual stimulus presentations and were less frequent after repeated visual stimulation. To better understand the role of 3-5 Hz oscillations in visual processing, we decided to investigate if 3-5 Hz oscillation prevalence and timing were affected by behavior in animals engaged in a visually guided decision making task.

### 3-5 Hz Vm oscillations occur during visual stimuli when animals performed a visually guided go/no-go task

To test if behavior modulated 3-5 Hz Vm oscillations, mice (n=17) were trained to perform a visually guided go/no-go discrimination task prior to whole-cell recordings (Figure 5A, Figure 5—figure supplement 1). During the task, animals had to decide whether to lick for a water drop (go) or withhold licking (no-go) based on visual cues ( go stimulus: 45° drifting gratings, no-go stimulus: 135° drifting gratings; Figure 5—figure supplement 1A). Visual stimuli were displayed for 3 seconds, and animals had to make their decision in the final second of the visual stimulus presentation (the response period). Animals reliably learned how to perform this task in 5 to 10 training sessions (Figure 5—supplement figure 1B). During training, animals’ licking behavior changed, especially, for go trials, where animals gradually began initiating licking prior to the response period (Figure 5—supplement figure 1C).

**Figure 5.**
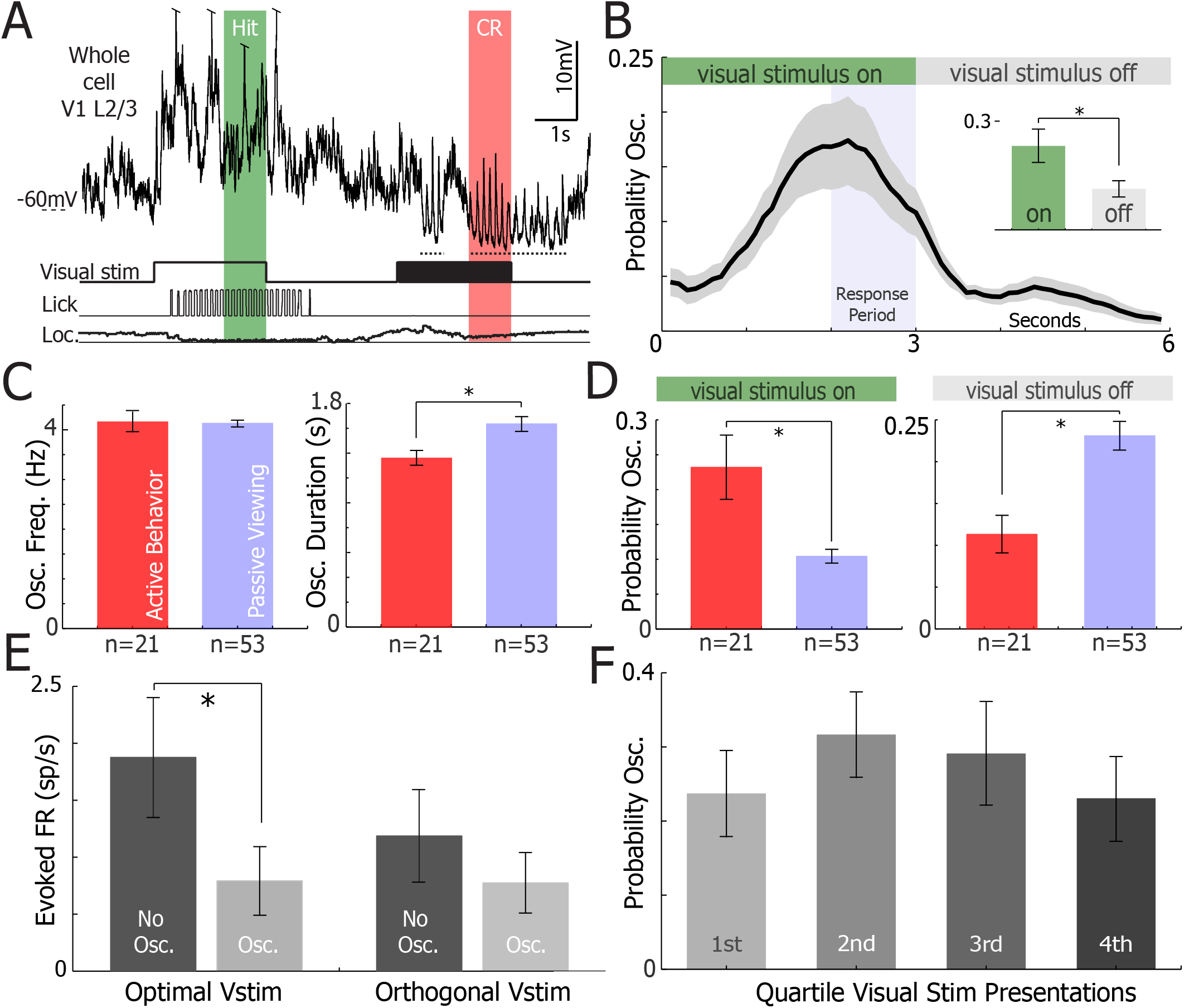
3-5 Hz oscillations occur predominately during visual stimulation while animals perform a visual discrimination task. (A) Example sub-threshold activity from a single neuron as animals performed the task. Visual stimuli timing, licking, and locomotion were recorded simultaneously. Arrows indicate instances of 3-5 Hz oscillations in the whole-cell recording. (B) The mean probability of 3-5 Hz oscillations occurring during a trial of the go/no-go task (n=21 neurons). Periods where visual stimuli were on and off are marked at the top. The response time, when the animal must report its decision, is denoted in the blue region. Shaded regions indicate ±SEM. Inset: the probability of an oscillation occurring when a visual stimulus was on and off. In contrast to passive viewing, 3-5 Hz oscillations occurred more frequently during visual stimuli presentations than during inter-trial intervals (WSRT, p = 0.026). (C) Comparison of the mean 3-5 Hz oscillation frequency (left, WRST, p = 0.8) and duration (right, WRST, p = 0.009) in neurons recorded from animals d uring active behavior (red, n=21) and passive viewing (blue, n=53). (D) Comparison of the mean probability of 3-5 Hz oscillations occurring in neurons recorded from animals during active behavior (red, n=21) and passive viewing (blue, n=53) while a visual stimulus is on (left, WRST, p = 0.001) and off (right, WRST, p = 0.007). (E) The mean firing rate evoked by optimal visual stimuli (left, WSRT, p = 0.001) and orthogonal visual stimuli (right, WSRT, p = 0.68) when 3-5 Hz oscillations were present ( osc.) or absent (no osc.) in neurons recorded from animals during active behavior (n=21). Error bars represent ±SEM. (F) The mean probability of 3-5 Hz oscillations occurrences was calculated during blocks of visual stimuli presentations grouped by the quartile of visual stimulus presentations. Neurons with less than 100 stimuli were excluded ( n=15, mean number of visual stimuli per neuron = 127 ± 14). No change in probability of 3-5 Hz oscillations was observed over the course of visual stimulus presentations (repeated measures one-way ANOVA, p = 0.099). Error bars represent ±SEM.

During active behavior, the onset time of 3-5 Hz oscillations was significantly different than during passive viewing and occurred almost exclusively during visual stimulus presentations (Figure 5B-D). Oscillations were initiated on average 1.71 ± 0.12 seconds (n = 21 neurons) after visual stimulus onset and were twice as likely to occur during visual stimulation than during inter-trial intervals (n=21 neurons, WSRT, p = 0.026; Figure 5B inset). As a result, 3-5 Hz oscillation probability during visual stimulation was significantly greater during active behavior than during passive viewing (WRST, p = 0.001; Figure 5D left). In contrast, 3-5 Hz oscillation probability following visual stimulation was significantly greater during passive viewing than during active behavior (WRST, p = 0.007; Figure 5D, right). The duration of oscillation epochs was slightly longer during active behavior compared to passive viewing (WRST, p < 0.009), but oscillation frequency was unchanged (WRST, p = 0.8; Figure 5C). Locomotion did not change oscillation prevalence or duration during active behavior (n=21, WSRT, p = 0.76 and p = 0.56, respectively; Figure 5— figure supplement 2A, B). As the go and no-go visual stimuli differed by 90°, one visual stimulus (the optimal visual stimulus) typically evoked a larger response than the other (the orthogonal visual stimulus) (Figure 5E). 3-5 Hz oscillations significantly reduced visually evoked action potential firing during optimal visual stimulus presentations (WSRT, p = 0.001), but not during the orthogonal visual stimulus presentations (n=21, WSRT, p = 0.68; Figure 5E). In contrast to passive viewing, the prevalence of oscillations did not decrease across the behavioral sessions (repeated measures one-way ANOVA, p = 0.099; Figure 5F). Therefore, oscillations reduced neuronal responsiveness to preferred visual stimuli during active behavior. These findings support the hypothesis that behavioral state plays a major role in modulating 3-5 Hz Vm oscillation prevalence and timing in V1.

### 3-5 Hz Vm oscillations’ prevalence and duration are modulated by behavioral response

3-5 Hz Vm oscillation prevalence and timing were also investigated in the context of animals’ responses during visually-guided behavior (Figure 6). 3-5 Hz oscillation prevalence was significantly higher during trials when animals correctly withheld licking (correct rejection, CR) than during trials when animals initiated a licking response either correctly (hit) or incorrectly (false-alarm, FA) (n=21, WSRT Bonferroni Corrected p = 0.046, p = 0.04, respectively). Importantly, the visual stimulus was identical in FA and CR trials, showing that behavioral response alone and not the sensory stimulus modulated oscillation prevalence. Yet, there was no difference in oscillation prevalence between incorrect and correct behavioral response (Hit vs FA, WSRT Bonferroni Corrected, p = 0.9; CR vs Miss, WSRT Bonferroni Corrected, p = .86). Additionally, oscillation duration was slightly longer during CR trials than during hit trials (WSRT Bonferroni Corrected, p = 0.035), but not FA trials (WSRT Bonferroni Corrected, p = 0.3). The high prevalence of oscillations during CR trials disprove the hypothesis that the motor action associated with licking response triggers oscillations because licking is typically absent during CR trials. Moreover, animals did not receive rewards during CR trials, indicating that reward expectation was not the primary factor in evoking 3-5 Hz oscillations in V1.

**Figure 6.**
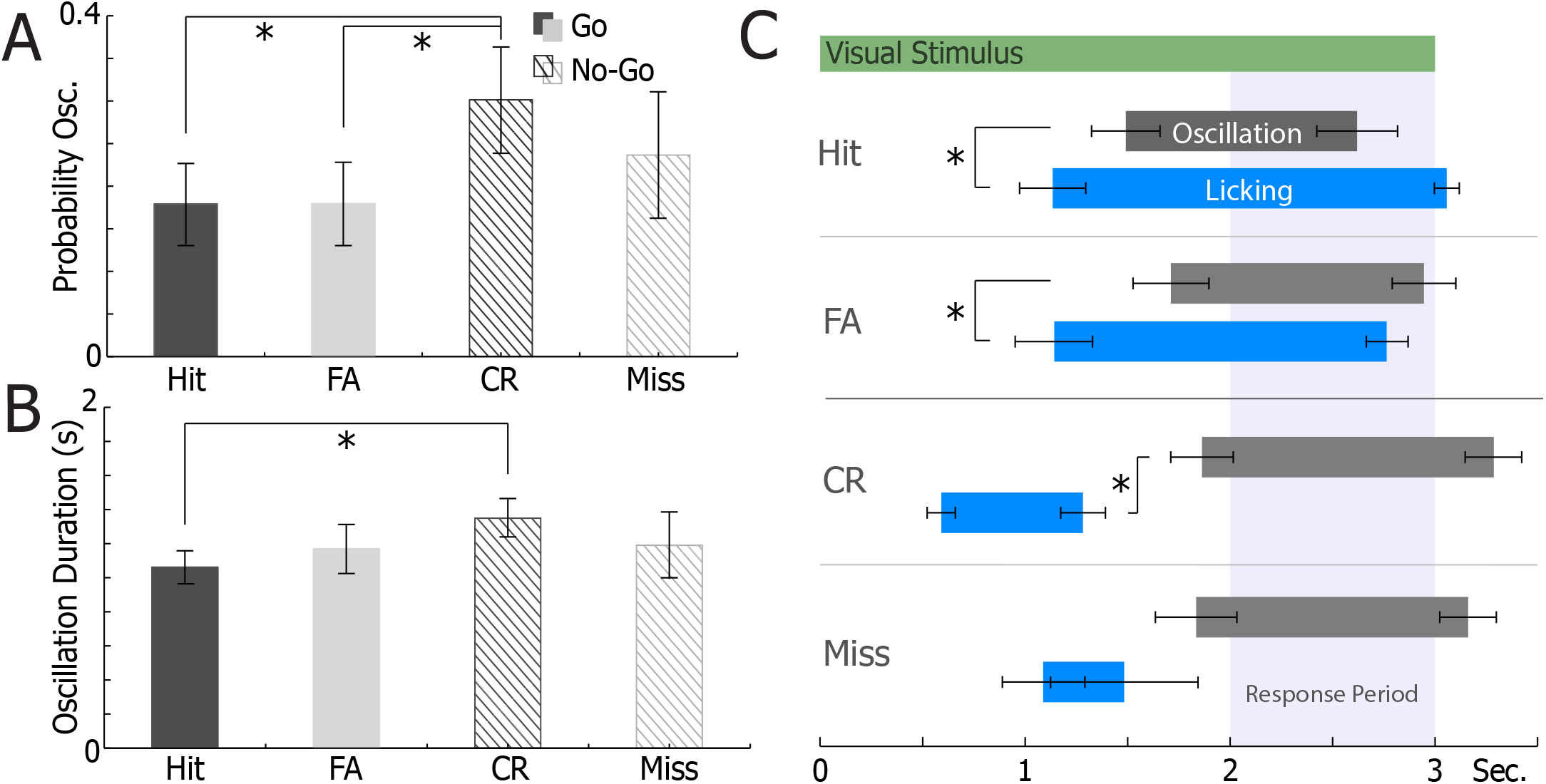
Behavioral response modulates oscillation probability and timing. (A) The mean probability of 3-5 Hz oscillations occurring during go trials (hits, black; false alarms (FA)) and no-go trials (correct rejections (CR), dark lines; misses, light lines) (n=21 neurons). Compared to CR trials, oscillations were less likely to occur during hit trials (WSRT Bonferroni corrected, p = 0.046) and FA trials (WSRT Bonferroni Corrected, p = 0.04). Error bars represent ±SEM. (B) The mean duration of 3-5 Hz oscillations during go trials ( hits, black; false alarms (FA)) and no-go trials (correct rejections (CR), dark lines; misses, light; n=21 neurons). Oscillations were shorter during hit trials than during CR trials (WRST Bonferroni Corrected, p = 0.035). Error bars represent ±SEM. (C) Comparison of oscillation (dark grey) and licking (blue) timing during h it, FA, CR and miss trials (n=21 neurons). Oscillations tend to begin after li cking onset in hit (WSRT, p = 0.01) and FA (WSRT,p = 0.001) trials. In CR trials with premature licking, oscillations tend to begi n after licking offset (WSRT, p = 0.031). Visual stimulus on time is indicated at the top. The response time is indicated in the light blue box. Error bars represent ±SEM.

3-5 Hz oscillation onset occurred after licking onset for correct(Hit, WSRT, p = 0.01) and incorrect(FA, WSRT, p = 0.001) go responses (Figure 6C). For trials where licking preceded the response period in correct no-go trials (CR), licking offset occurred prior to 3-5 Hz oscillation onset (WSRT, p = 0.031). There was no difference in oscillation onset time across behavioral responses (Repeated Measures one-way ANOVA, p = 0.35). Therefore, 3-5 Hz oscillations followed the animal’s response to the go/no-go visual cue.

### 3-5 Hz oscillations are absent from V1 L2/3 neurons when animals perform an analogous auditory decision making task

To test whether 3-5 Hz oscillations in V1 were specific to processing of visual information during visual discrimination, V1 neurons’ Vm was recorded as animals performed an analogous auditory go/no-go task (Figure 7). All task parameters were identical with the exception that animals based their decision on auditory cues (5 kHz – go, 10 kHz – no-go; Figure 7A) and no visual stimuli were shown. During the auditory task, a monitor was placed in the identical position as during the visual tas k, and an isoluminant grey screen was displayed throughout the recording to provide equal illumination as during the visual task. Oscillations occurred much less frequently when animals based their decision on auditory cues instead of visual cues (Figure 7B, C, & D). The probability of a 3-5 Hz oscillation occurring during stimulus presentations increased approximately four-fold during the visual task than the auditory task (auditory n = 7, visual n= 21, p = 0.003 WRST). Yet, no difference was detected in oscillation duration (WRST, p = 0.27) and oscillation onset latency from stimulus onset (WRST, p = 0.64) between animals performing the visual and auditory tasks (Figure 7D). Finally, animals discriminated between auditory and visual stimuli equally well (WRST, p = 0.37), indicating that animal performance was not different during visual and auditory tasks. Taken together, these results suggest that 3-5 Hz oscillations in V1 neurons were primarily associated with visual information processing as opposed non-specific decision making and motor outputs associated with the task.

**Figure 7.**
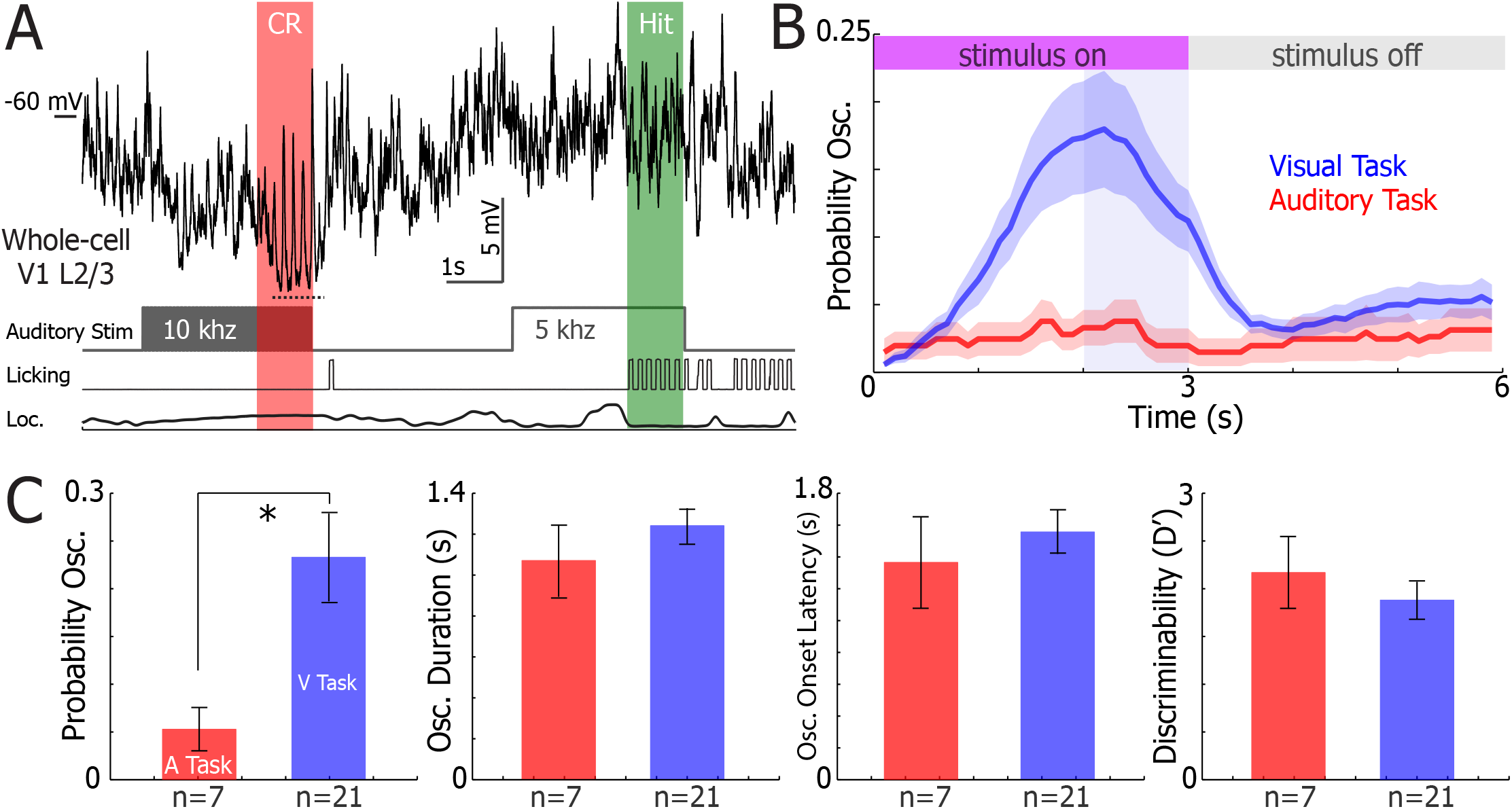
3-5 Hz oscillations are absent in V1 when animals perform an analogous auditory discrimination task. (A) Example sub-threshold activity from a single neuron as animals performed the task. Auditory stimuli timing, licking, and locomotion were recorded simultaneously. Arrow indicates an instance of 3-5 Hz oscillations in the whole-cell recording. (B) The mean probability of 3-5 Hz oscillations occurring during a trial of the auditory (n=7 neurons) and visual (n=21 neurons) go/no-go tasks. Periods where stimuli were on and off are marked at the top. The response time, when the animal must report its decision, is denoted in the light blue region. Shaded regions indicate ±SEM. (C) Comparison between the mean probability of 3-5 Hz oscillations during a trial (WRST, p = 0.003), oscillation duration (WRST, p = 0.27), oscillation onset latency from stimulus onset (WRST, p = 0.64), and discriminability (WRST, p = 0.037) during the auditory (red) and visual (blue) discrimination task. Error bars represent ±SEM.

## DISCUSSION

We performed two-photon guided whole-cell recordings in awake mice to investigate a novel gain reduction mechanism in L2/3 V1 neurons of mice. We found that 3-5 Hz subthreshold oscillations decreased the gain of excitatory neurons but not PV+ and SOM+ interneurons, which oscillated in phase with excitatory neurons and fired strongly at the depolarized peaks of oscillations. In addition, oscillation recruitment relied both on visual processing and the animal’s behavioral state. As a result, 3-5 Hz subthreshold oscillations represent a gain reduction mechanism which adjusts neuronal activity according to an animal’s sensory and behavioral context.

3-5 Hz subthreshold oscillations may decrease the gain of excitatory neurons to sensory cues in at least one of the following ways: (a) the hyperpolarized Vm baseline during oscillatory sequences likely contributes to decrease the gain of the neurons by reducing the response magnitude to incoming signals (Cardin et al., 2008; Carandini & Ferster, 1997; Nowak et al., 2005); (b) during the depolarizing phases of the oscillations where excitatory neurons’ Vm is closest to reaching spike threshold, excitatory neurons received strong perisomatic and dendritic inhibition from GABAergic PV+ and SOM+neurons, respectively (Taniguchi, 2014; Figure 2); (c) sensory signals out of phase with 3-5 Hz oscillations could filter inbound sensory signals of a different time structure (Engel et al., 2001; Schroeder & Lakatos, 2009; Lakatos et al., 2008). Considering the combination of these three mechanisms, 3-5 Hz subthreshold oscillations represent a potent combination of inhibitory strategies to reduce the gain of excitatory sensory neurons.

3-5 Hz subthreshold oscillations may be important in other cortical circuits as they have been observed in barrel (Poulet & Petersen, 2008), auditory (Zhou et al., 2014; Schneider et al., 2014) and motor (Zagha et al., 2015) cortex neurons in awake behaving mice. In particular, Zagha and colleagues investigated the distribution of the Vm of M1 neurons during 3-5 Hz subthreshold oscillations and observed an approximate 8 mV hyperpolarization of the mean membrane potential, which reduced the probability that the M1 neuron’s Vm would cross the spike threshold. Simultaneous with the subthreshold oscillations, 3-8 Hz LFP power was significantly higher in S1 and M1 during miss trials while animals performed a whisker deflection detection task. Zagha and colleagues hypothesized that these oscillations disorganized task-relevant circuitry by correlating activity in opposing neural ensembles. As a result, 3-5 Hz subthreshold oscillations likely exist beyond the visual cortex and could perform a similar function in other sensory cortices.

The behavioral significance of 3-5 Hz subthreshold oscillations in visual cortex may be to reduce processing of behaviorally irrelevant visual stimuli. Accordingly, we found that oscillations were most prevalent after animals had made their decision during visual discrimination (Figure 6), a point in the task when additional visual inputs were irrelevant to completing the task. This finding alone would predict that oscillations would occur whenever animals do not require visual input during decision making, such as when animals perform an auditory discrimination task. Instead, we found that 3-5 Hz oscillations were not evoked when animals did not engage in visual cues (Figure 7), illustrating that engagement with visual stimuli is critical for eliciting oscillations. In fact, the level of animal engagement with visual stimuli may influence the prevalence of oscillations given that oscillation prevalence decreased over time during passive viewing (Figure 4C) but not during active visual discrimination (Figure 5G). Therefore, we propose that 3-5 Hz subthreshold oscillations may be evoked during visual information processing to decrease the gain of V1 neurons at times when visual cues are no longer behaviorally relevant.

Such a mechanism could be particularly useful during other behaviors such as attention and working memory. When non-human primates ignore visual cues during attention tasks, neurons in V4 increase their correlated firing at frequencies between 3 and 5 Hz, spiking synchronizes within low frequency bands (<10 Hz) of the LFP (Mitchell et al., 2009; Fries et al., 2001), and LFP power between 3-5 Hz increases (Fries et al., 2008). During visually-guided working memory tasks in non-human primates, prominent high-amplitude 4-8 Hz LFP oscillations appear in visual cortex and synchronize single-unit firing to the peaks of the oscillations during the delay period (Lee et al., 2005; Liebe et al., 2012). If coordinated subthreshold oscillations are responsible for producing these LFP and spiking patterns, their role may be to exclude processing of unattended cues during attention and task irrelevant visual information during working memory.

3-5 Hz oscillation generation could be the result of resonant activity in the thalamocortical network. The thalamocortical loop is responsible for generating several natural and pathological oscillations, including oscillations in the 3-5 Hz range (Steriade et al., 1993, Destexhe & Sejnowski, 2003, Buzsáki & Draughn, 2004). Thalamocortical neurons switch between tonic spiking and oscillatory burst firing depending on their resting membrane potential, a phenomenon largely due to low-voltage activated T-type Ca^2+^ channels (Jahnsen & Llinás, 1984; Contreras, 2006; Halassa, 2012). Neuromodulatory inputs, including cholinergic and monoaminergic sources, regulate the resting membrane potential of thalamic neurons to allow or block the generation of oscillations (McCormick, 1989; Saper et al., 2005; Steriade et al., 1993). Given that neuromodulatory tone can play a key role in modulating visual processing (Polack et al., 2013; Pinto et al., 2013; McCormick et al., 1993; Disney et al., 2007; Chubykin et al., 2013), it is conceivable that 3-5 Hz oscillations could be caused by a change in thalamic neuromodulation, allowing thalamocortical neurons to hyperpolarize and enter a burst state capable of generating 3-5 Hz oscillations.

In conclusion, it is possible that the mechanism identified in this study may modulate cortical computations in a variety of cortical circuits during several different behaviors. More work will be needed to fully understand the cellular and network properties and functional significance of subthreshold 3-5 Hz oscillations. In particular, further studies will focus on understanding how and where these oscillations are generated. Finally, it will be important to record subthreshold oscillations in other brain areas during different behavioral tasks to confirm whether this mechanism is indeed ubiquitous in cortical circuits.

## MATERIALS AND METHODS

### Surgery

All experimental procedures were approved by the University of California, Los Angeles Office for Animal Research Oversight and by the Chancellor’s Animal Research Committees. Adult (2–12 months old) male and female C57Bl6/J, SOM-Cre (JAX number 013044) × Ai9 (JAX number 007909), and PV-Cre (JAX number 008069) × Ai9 mice were anesthetized with isoflurane (3–5% induction, 1.5% maintenance) ten minutes after injection of a systemic analgesic (carprofen, 5 mg per kg of body weight) and placed in a stereotaxic frame. Mice were kept at 37°C at all times using a feedback-controlled heating pad. Pressure points and incision sites were injected with lidocaine (2%), and eyes were protected from desiccation using artificial tear ointment. The skin above the skull was incised, a custom-made lightweight metal head holder was implanted on the skull using Vetbond (3M) and a recording chamber was built using dental cement (Ortho-Jet, Lang). Mice had a recovery period from surgery of five days, during which they were administered amoxicillin (0.25 mg per ml in drinking water through the water supply). After the recovery period, mice were habituated to head fixation on the spherical treadmill. On the day of the recording, mice were anesthetized with isoflurane. To fix the ground wire, a small craniotomy (.5 mm diameter) was made above the right cerebellum and a silver wire was implanted at the surface of the craniotomy and fixed with dental cement. A circular craniotomy (diameter = 3 mm) was performed above V1 and a 3-mm diameter coverslip drilled with a 500-μm diameter hole was placed over the dura, such that the coverslip fit entirely in the craniotomy and was flush with the skull surface. The coverslip was kept in place using Vetbond and dental cement, and the recording chamber was filled with cortex buffer containing 135 mM NaCl, 5 mM KCl, 5 mM HEPES, 1.8 mM CaCl2 and 1 mM MgCl2. The head-bar was fixed to a post and the mouse was placed on the spherical treadmill to recover from anesthesia. All recordings were performed at least two hours after the end of anesthesia, when the mouse was alert and could actively participate in the behavioral task.

### Electrophysiological recordings

Long-tapered micropipettes made of borosilicate glass (1.5-mm outer diameter, 0.86-mm inner diameter, Sutter Instrument) were pulled on Sutter Instruments P-1000 pipette puller to a resistance of 3–7 MΩ, and filled with an internal solution containing 115 mM potassium gluconate, 20 mM KCl, 10 mM HEPES, 10 mM phosphocreatine, 14 mM ATP-Mg, 0.3 mM GTP, and 0.01–0.05 mM Alexa-594 (for experiments with C57Bl/6 mice) or Alexa-488 (for interneuron recordings). Pipettes were lowered into the brain under two-photon imaging guidance performed with a Sutter MOM microscope using a Ti - Sapphire Ultra-2 laser (Coherent) at 800 nm and a 40× 0.8 NA Olympus water-immersion objective. Images were acquired using Scanimage 3.2 software (Pologruto et al., 2003) Whole-cell current-clamp recordings were performed using the bridge mode of an Axoclamp 2A amplifier (Molecular Devices), then further amplified and low-pass filtered at 5 kHz using a Warner Instruments amplifier (LPF 202A). Recordings typically lasted 30 min (range 5 to 50 min). Recordings or parts of recordings with unstable membrane potential and/or action potentials < 35 mV were excluded from analysis. ECoG recordings were performed with an alternating/direct current differential amplifier (Model 3000, A-M system) and band-pass filtered at 0.1–3,000 Hz. Analog signals were digitized at 12 kHz with WinEDR (Strathclyde University) using a NIDAQ card (National Instruments). We recorded 40 excitatory, 6 PV+, and 7 SOM+ neurons from 29, 5, and 6 untrained mice, respectively, in separate experiments to ascertain 3-5 Hz oscillation activity during spontaneous behavior and passive viewing. We recorded 21 neurons from 17 trained mice in separate experiments to ascertain 3-5 Hz oscillation activity during visual and auditory discrimination.

### Visual Stimulus Presentation

A 40-cm diagonal LCD monitor was placed in the monocular visual field of the mouse at a distance of 30 cm, contralateral to the craniotomy. Custom-made software developed with Psychtoolbox in MATLAB was used to display drifting sine wave gratings (series of 12 orientations spaced by 30 degrees randomly permuted, temporal frequency = 2 Hz, spatial frequency = 0.04 cycle per degree, contrast = 100%). For passive viewing, the presentation of each orientation lasted 1.5 or 3 s and was followed by the presentation of a gray isoluminant screen for an additional 1.5 or 3 s, respectively. The electrophysiological signal was digitized simultaneously with two analog signals coding for the spatial and temporal properties of the grating. The treadmill motion was measured every 25 ms (40 Hz) by an optical mouse whose signal was converted into two servo pulse analog signals (front-back and left-right) using an external PIC microcontroller, and acquired simultaneously with the electrophysiological data.

### Training

C57Bl/6J mice (Jackson Labs) with head-bar implants were water-deprived to 90% of their body weight and acclimated to head-fixation on a spherical treadmill in custom-built, sound-proof training rigs. Each rig was equipped with a monitor (Dell), water dispenser with a built-in lickometer (to monitor licking, infrared beam break) (Island-Motion), an infrared camera (Microsoft), and stereo speakers (Logitech). In addition, data acquisition boards (National Instruments) were used to actuate water delivery and vacuum reward retrieval as well as monitor animal licking. Data acquisition boards and the monitor were connected to a laptop (Dell), which ran the custom made training program (MATLAB). Once animals reached the target weight, they were trained to discriminate visual stimuli or auditory. In the visual discrimination task, drifting sine-wave gratings at one orientation were paired with a water reward, and the animal was expected to lick ( go). Orthogonal drifting gratings signaled the absence of reward, and the animal was expected to withhold licking ( no-go) during these trials. In the auditory discrimination task, a 100 dB 5 kHz pure tone indicated Go trials and a 100 dB 10 kHz pure tone indicated No-Go trials.

Each trial lasted three seconds. The visual or auditory stimulus was present for the duration of the trial. When the stimulus instructed the animal to lick, water was dispensed two seconds after stimulus onset. No water was dispensed in the no-lick condition. Licking was only assessed during the final second of the trial. If the animal responded correctly, the inter-trial interval (ITI) was 3 seconds. If the animal responded incorrectly, the ITI was increased to 9.5 seconds as negative reinforcement. If the animal missed a reward, the reward was removed by vacuum at the end of the trial. Animals performed 300-500 trials daily.

Performance was measured using the D’ statistic (D’=norminv(fraction trials with correct licking) – norminv(fraction trials with incorrect licking), norminv = inverse of the normal cumulative distribution function), which compares the standard deviation from chance performance during lick and no-lick trials (chance D’=0). Animals were considered experts if their sessions average D’ > 1.7 (probability of chance behavior < 0.1%, Monte Carlo Simulation).

### Analysis

Data analysis was performed using custom made routines in MATLAB. The 3-5 Hz oscillations were defined as regular low frequency and high-amplitude oscillations of the Vm superimposed on a steady hyperpolarizing envelope (see examples in Figs. 1b, 3a, 3b, 5a, and 7a). The Vm baseline was defined as the mean of the bottom 20^th^ percentile of the Vm distribution, and the change in Vm baseline during oscillations was defined as the baseline during the oscillation epoch minus the baseline one second prior to the oscillation epoch. The spontaneous firing rate during the oscillation was calculated as the total number of action potential recorded during the oscillation divided by the duration of the oscillation. This was then compared to the firing rate measured during the 1.5 seconds preceding the oscillation. Phase offset was obtained by calculating the difference in time between positive peaks in low pass filtered (-3 dB @ 10 Hz) ECoG and Vm signals measured in degrees during oscillatory epochs.

The orientation selectivity index (OSI) in excitatory neurons was calculated using the following equation (Mazurek et al., 2014):*osi* = 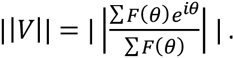 To compare firing rates evoked by visual stimuli during passive viewing and behavior, trials with the presence of an oscillatory epoch at any point of the trial were compared to trials without any oscillations. Oscillation incidence was defined as the number of oscillations occurring over all spontaneous activity or passive viewing divided by the total time. Probability of oscillation and oscillation onset was defined as the probability of the event occurring in a given time bin. During the behavioral task, the optimal visual stimulus was defined as the stimulus that had a greater mean evoked firing rate.

### Statistics

Unless stated otherwise, statistical significance was calculated by Wilcoxon Signed Rank Test (WSRT), Wilcoxon Rank-Sum Test (WRST), One Way Analysis of Variance (ANOVA), and Repeated Measures one-way ANOVA. Scale bars and shading around means represent SEM unless indicated. Wilcoxon tests were performed in MATLAB and ANOVA tests were performed in SPSS Statistics version 21 (IBM).

## ACKNOWLEDGEMENTS

The authors declare no competing financial interest. We thank all the members of the Golshani Lab for their support and insightful comments in reviewing the data and manuscript. This work was supported by NIH grants 1R01-MH101198-01 and R01-MH105427-A1. M.E. is supported by a National Research Service Award F31EY025185-02. P-O. P. performed the electrophysiological recordings in non-behaving mice. M.E. and P-O.P. performed the electrophysiological recordings in behaving mice. M.E., P-O.P., and P.G. designed the study. M.E. and P-O.P. designed the behavioral paradigm. M.E. analyzed the data M.E. wrote the manuscript with contribution from P.G. and P-O.P. Correspondence and requests for materials should be addressed to.

---

**Figure 3— figure supplement 1.**
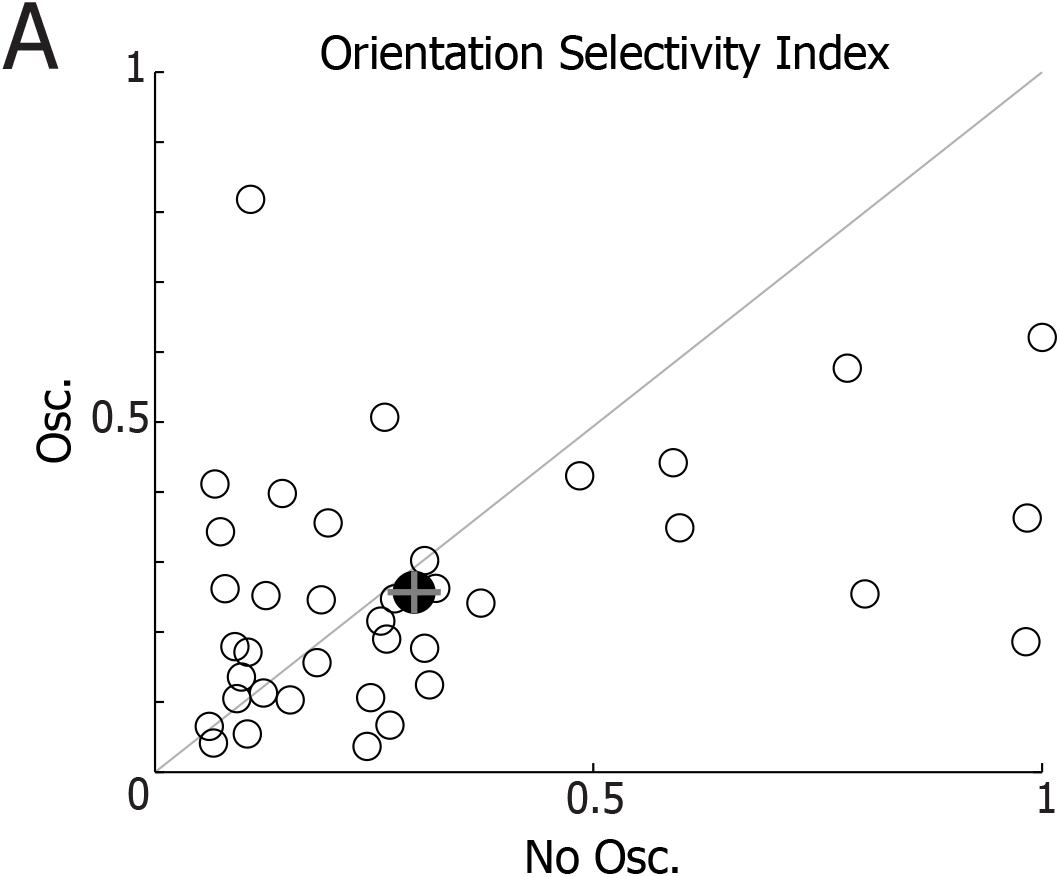
Oscillations do not affect the orientation selectivity index of excitatory neurons. (A) The orientation selectivity index of excitatory neurons was calculated for excitatory neurons during passive viewing when 3-5 Hz Vm oscillations were present (osc.) or not present (no osc.; see methods for calculation). Orientation selectivity was not changed by the presence of 3-5 Hz Vm oscillations (n = 40; WSRT, p = 0.93).

**Figure 4—figure supplement 1.**
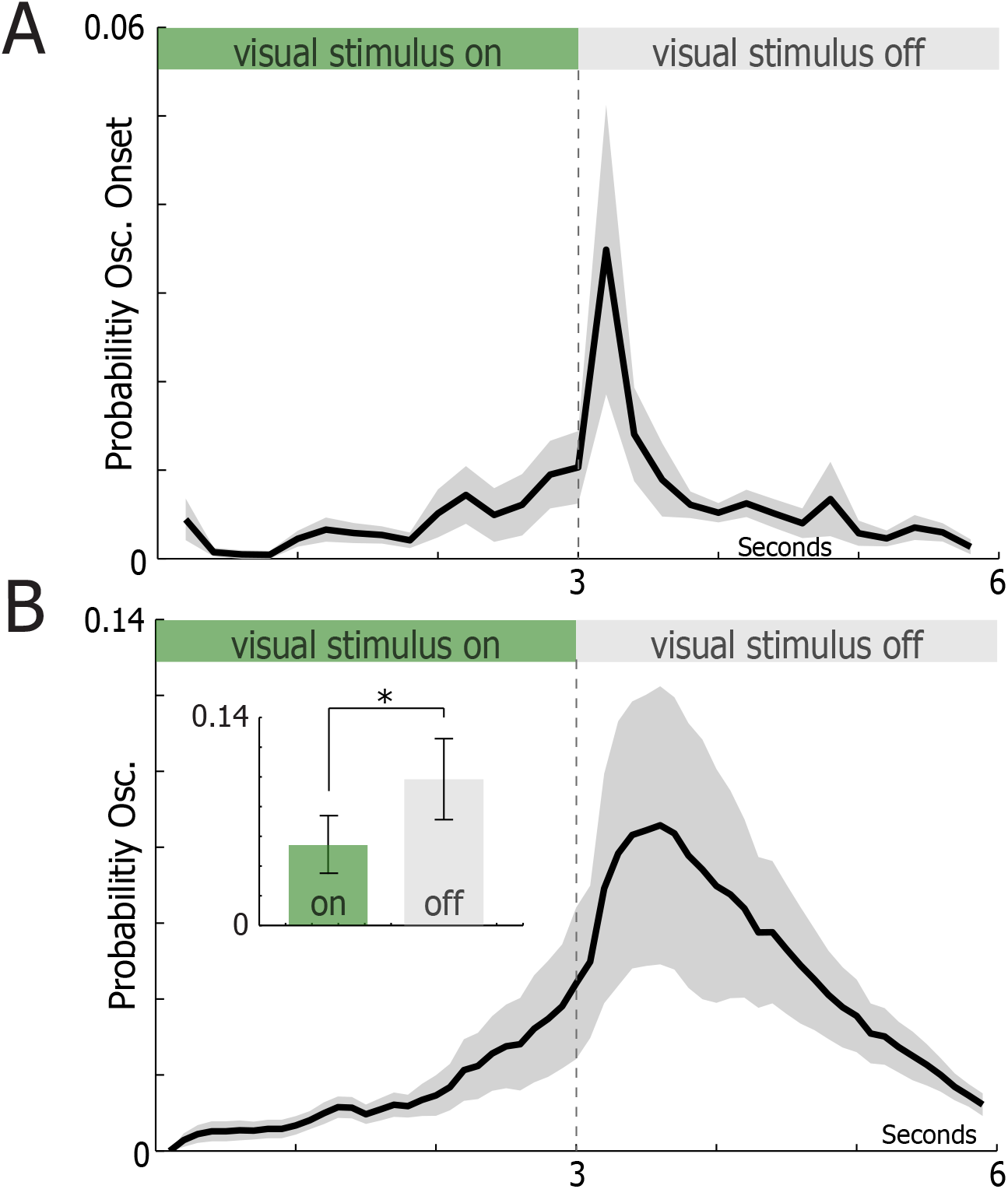
Oscillation timing is shifted proportionally when the visual stimulus duration is increased. (A) The mean probability of 3-5 Hz oscillation onset during and after drifting gratings presents for three seconds (n = 9 neurons). Shaded regions indicate ±SEM. (B) The mean probability of 3-5 Hz oscillations during and after drifting gratings presented for three seconds (n = 9 neurons). Shaded regions indicate ±SEM. Inset: the probability of an oscillation occurring when a visual stimulus was on and off. Oscillations occurred more frequently between visual stimuli presentations than during visual stimuli presentations (WSRT, p = 0.004).

**Figure 4—figure supplement 2.**
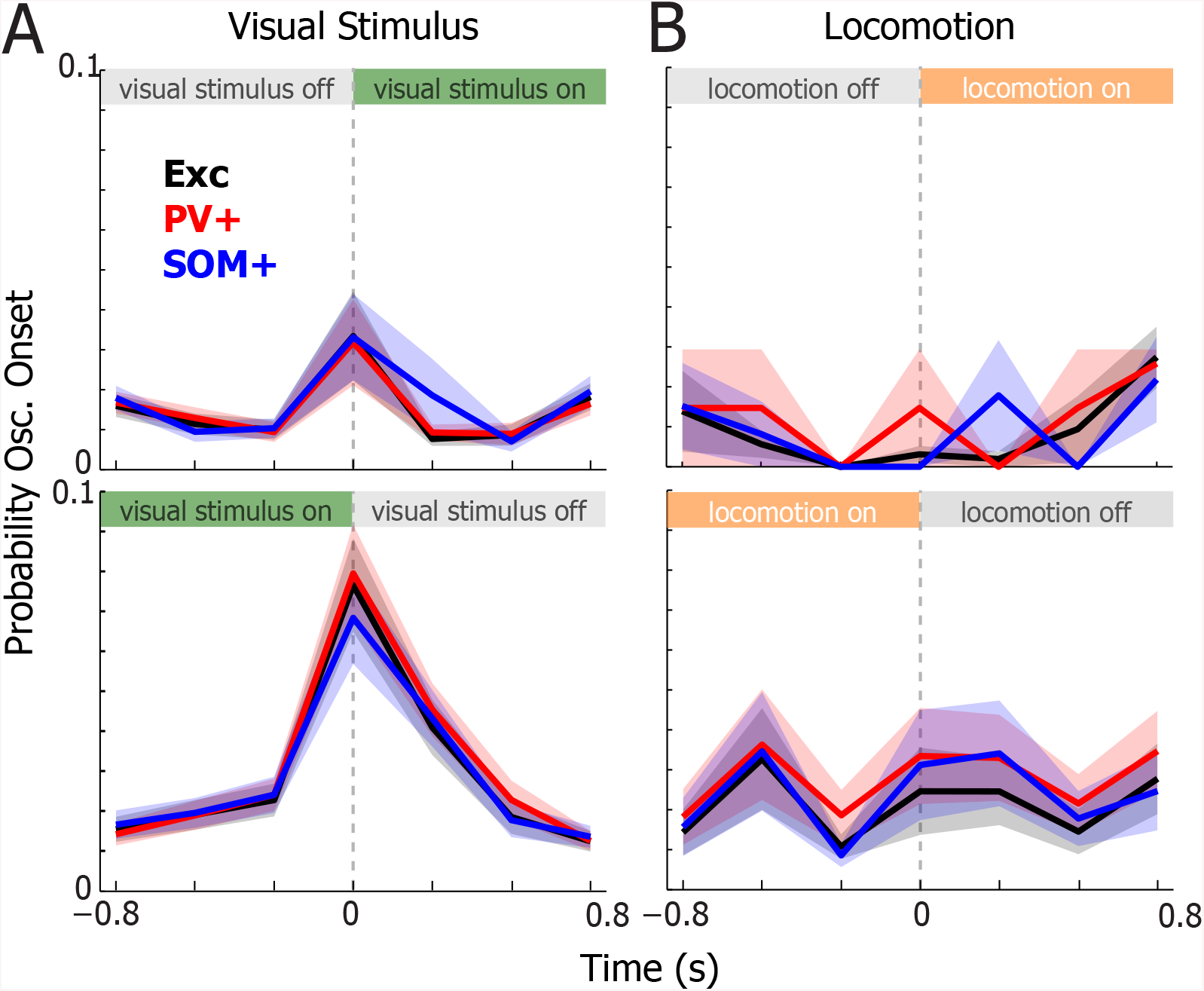
Probability of oscillation onset at visual stimulus and locomotion onset and offset. (A) The mean probability of 3-5 Hz oscillation onset at visual stimulus onset (top) and offset (bottom) for excitatory (black, n=40), PV+ (red, n=6), and SOM+ (blue, n=7) neurons. Shaded regions indicate ±SEM.(B) The mean probability of 3-5 Hz oscillation onset at locomotion onset (top) and offset (bottom) for excitatory, PV+, and SOM+ neurons. Shaded regions indicate ±SEM.

**Figure 5— figure supplement 1.**
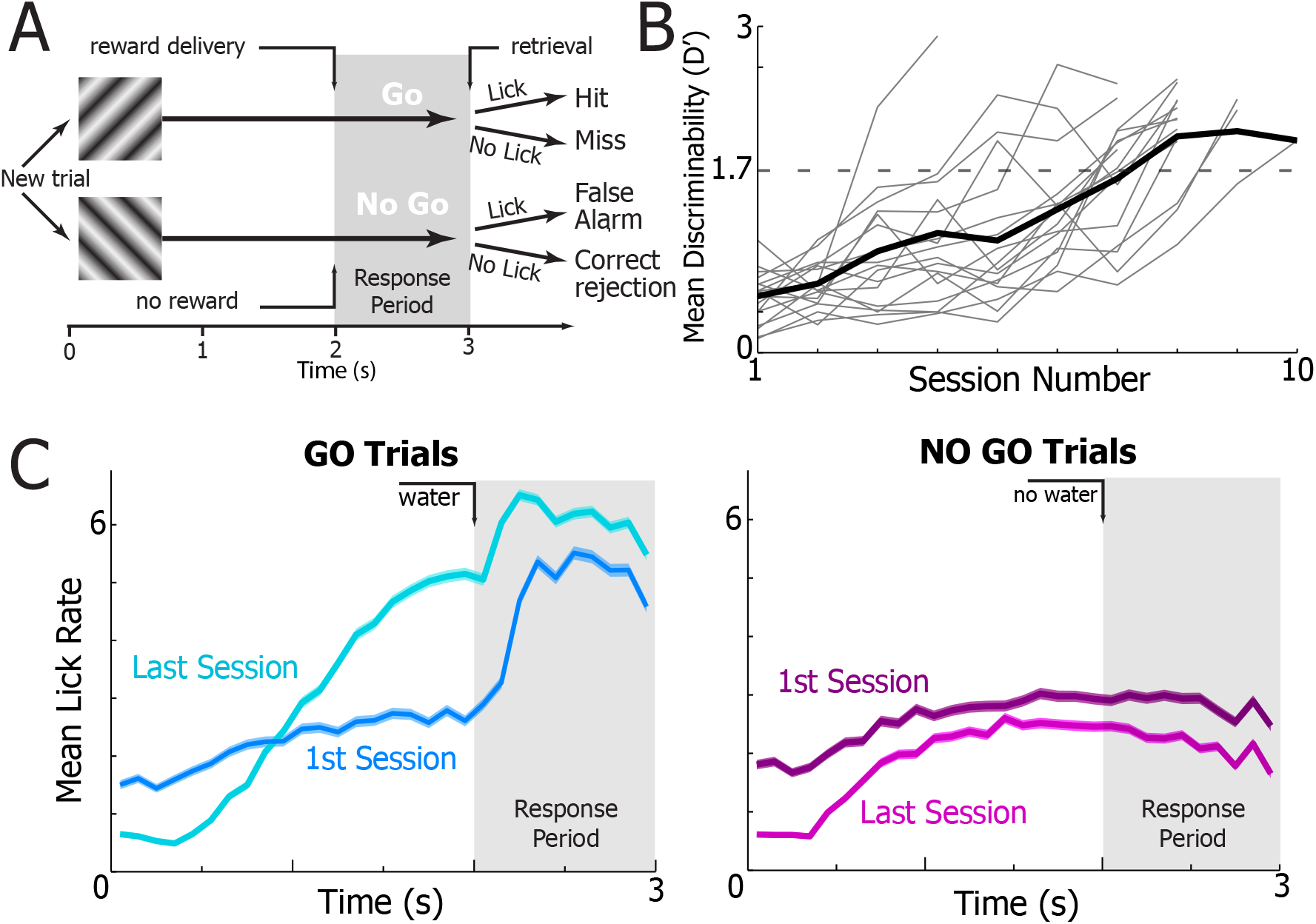
Task schematic and animal learning curves. (A) Left: Schematic of the training set-up. Right: Task schematic. Visual stimuli were presented for three seconds. In go trials, 45° gratings were displayed and a water reward was issued two seconds after stimulus onset. During no-go trials, 135° gratings were displayed and no reward was issued. Animal response (licking) was recorded during the response period to assess correct behavior. For more details, see Materials and Methods. (B) The mean discriminability of animals during training, which is a measure of animal performance (n = 17 mice). Black line: the mean performance of all animals on a given session date. Light grey lines: the mean performance of a single animal on a given session date. Animals were recorded once their mean discriminability surpassed D’=1.7 (Monte Carlo Simulation, p = 0.01 random behavior). (C) The mean lick rate of animals during go (left) and no-go (right) trials during their first training session (darker) and last session (lighter).

**Figure 5— figure supplement 2.**
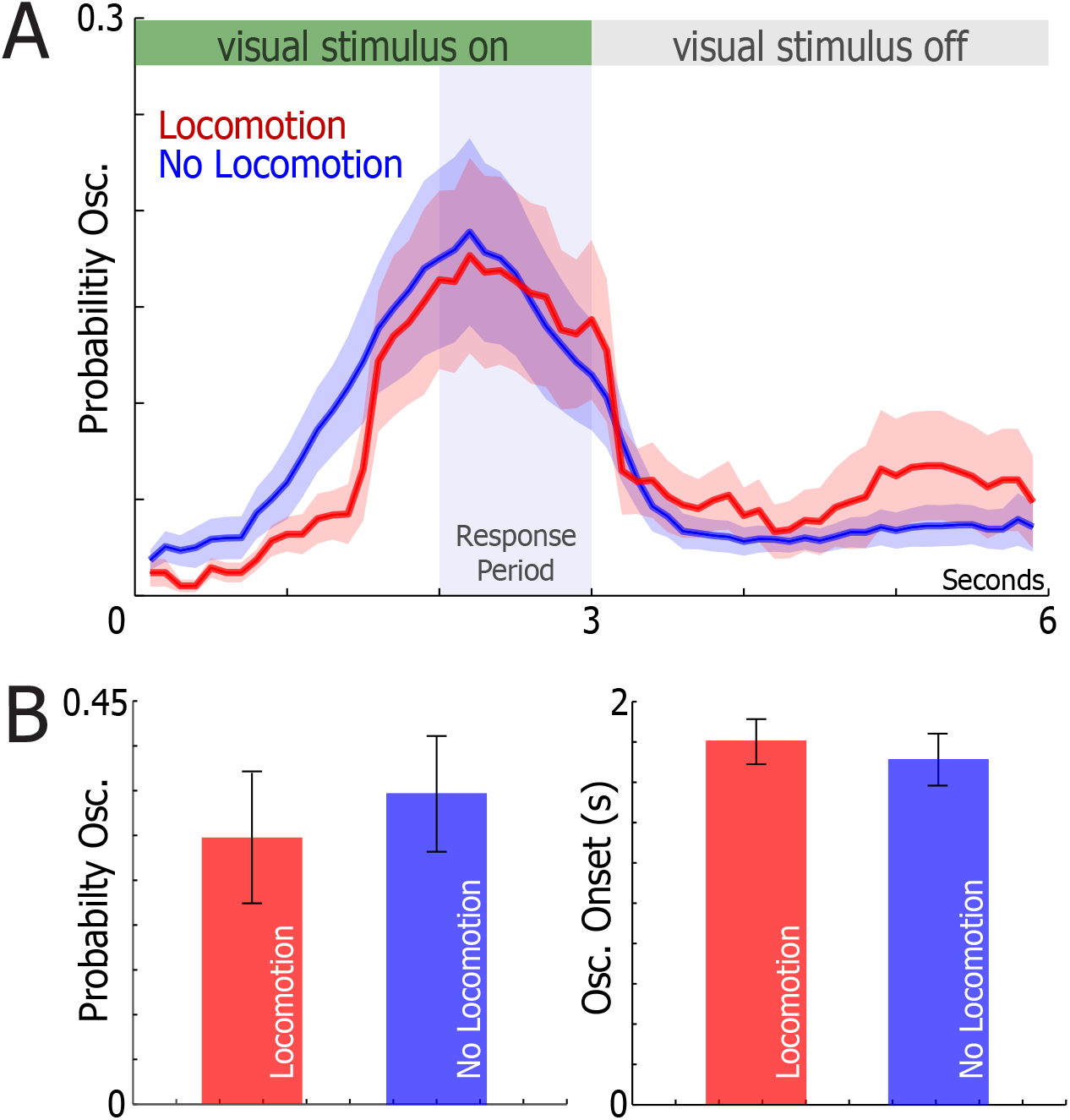
Locomotion does not change 3-5 Hz oscillation probability during active behavior. (A) The mean probability of 3-5 Hz oscillations occurring during trials of the visual discrimination task with locomotion (red) and without locomotion (blue)(n=21 neurons). Visual stimuli on and off times are shown at the top. The response time is indicated by the blue box. Shaded regions represent ±SEM. (B) The mean oscillation probability (left) and oscillation onset latency from visual stimulus onset (right) during trials with (red) and without (blue) locomotion (n=21 neurons). No changes in oscillation probability (WSRT, p = 0.76) and oscillation onset latency (WSRT, p = 0.56) were observed between trials with locomotion and without locomotion.

